# Frontoparietal connectivity correlates with working memory performance in multiple sclerosis

**DOI:** 10.1101/639930

**Authors:** Alejandra Figueroa-Vargas, Claudia Cárcamo, Rodrigo Henríquez-Ch, Francisco Zamorano, Ethel Ciampi, Reinaldo Uribe, Macarena Vásquez, Francisco Aboitiz, Pablo Billeke

## Abstract

Working Memory (WM) impairment is the most common cognitive deficit of Multiple Sclerosis (MS) patients. However, evidence of its neurobiological mechanisms is scarce. Here we recorded electroencephalographic activity of twenty patients with relapsing-remitting MS and minimal cognitive deficit, and 20 healthy control (HC) subjects while they solved a WM task. In spite of similar performance, the HC group demonstrated both a correlation between temporoparietal theta activity and memory load, and a correlation between medial frontal theta activity and successful memory performances. MS patients did not show theses correlations leading significant differences between groups. Moreover, cortical connectivity analyses using granger causality and phase-amplitude coupling between theta and gamma revealed that HC group, but not MS group, presented a load-modulated progression of the frontal-to-parietal connectivity. This connectivity correlated with working memory capacity in MS groups. This early alterations in the oscillatory dynamics underlaying working memory could be useful for plan therapeutic interventions

## Introduction

Multiple Sclerosis is a chronic demyelinating disease of the Central Nervous System^1^. The physiopathology is based on deregulation of the immune system generating motor, cognitive, and neuropsychiatric symptoms^2,3^. Cognitive impairments are present in 40 to 70% of patients, affecting their professional development, personal relationships, mood and quality of life^4,5^. These symptoms can be detected from early phases of the disease and may include alterations in information processing speed, attention, executive functions, and working memory (WM)^6,7^. Since human cognition critically depends on WM, an ability that enables us to adaptively maintain and manipulate information according to the demands of the environment, alterations in this process seem to be a key step in cognitive alterations in patients with multiple sclerosis. Researchers confirmed WM alterations are a habitual impairment, affecting patients early in the course of the disease^8,9^.

WM is a hierarchical process that links sensory representations to specific responses, through intermediate representations relevant to the task and action plans^10^. A distributed network of brain areas participates in this process, exhibiting sustained activity during the period of WM maintenance in the absence of sensory stimuli^11^. This sustained activity is generated by reverberant discharges in an interconnected network that involves the prefrontal cortex (PFC), the posterior parietal and temporal lobes^11^. Electrophysiological studies have revealed the crucial participation of oscillatory activity in theta (4-8 Hz), alpha (8-12 Hz), and gamma (30-100 Hz) bands in WM^12,13^. During the maintenance stage, memory load increases theta activity and theta-gamma coupling in frontal and temporo-parietal regions^14–16^. EEG and MEG studies have shown a greater synchrony between the frontal and parietal regions associated with the amount of information successfully maintained in WM^15,16^.

The neurobiological mechanism underlying WM alterations in multiple sclerosis are not well known. Successful WM needs the coordination of distributed brain networks that are especially sensitive to the diffuse damage of white and grey matter found in multiple sclerosis^17,18^. Several reports indicate very early alterations in cortico-cortical connectivity in multiple sclerosis^19,20^. Both functional and structural brain imaging techniques have revealed alterations in brain network connectivity patterns in patients with minimal or low cognitive disabilities^19^. An EEG study found a decrease in alpha and theta band coherence between the anterior and posterior electrodes, as well as between inter-hemispheric regions during rest. These alterations seem to be related to both cognitive deficits and subcortical lesion burden^21^. Thus, a functional marker of the cognitive alterations for early stages of the disease with minimal clinical manifestations would be relevant to address early cognitive rehabilitation. Additionally, the identification of the precise functional alterations in the oscillatory patterns opens the opportunity to plan specific interventions using, for example, non-invasive brain stimulation. Therefore, the aim of our study was to assess the neurophysiological underpinnings of alterations in the cortical circuits that support WM in multiple sclerosis. We hypothesized that WM impairments in patients with multiple sclerosis are due to an impairment in the maintenance of activity in the fronto-parietal network that is reflected in a reorganization of cortical oscillatory dynamics. Specifically, we predicted that i) multiple sclerosis alters the progressive increases of power of both theta and alpha oscillatory activity in relation to the increases of memory load; and that ii) fronto-parietal theta connectivity underlying successful memory information maintenance is impaired in patients with multiple sclerosis.

To address this issue, we recorded EEG activity of forty individuals. Twenty patients had relapsing-remitting multiple sclerosis with minimal clinical cognitive alterations and twenty healthy control subjects (see Table 1 for demographic data) solved working memory tasks (see Figure 1). Subjects had to memorize two, four or six consonants, generating three levels of working memory load (see more detail in methods section). Importantly, the identification of the precise neurophysiological dynamics underlying WM alteration in patients with multiple sclerosis could contribute to both early detection and development of specific cognitive rehabilitation interventions ^11,22^.

**Figure 1.**
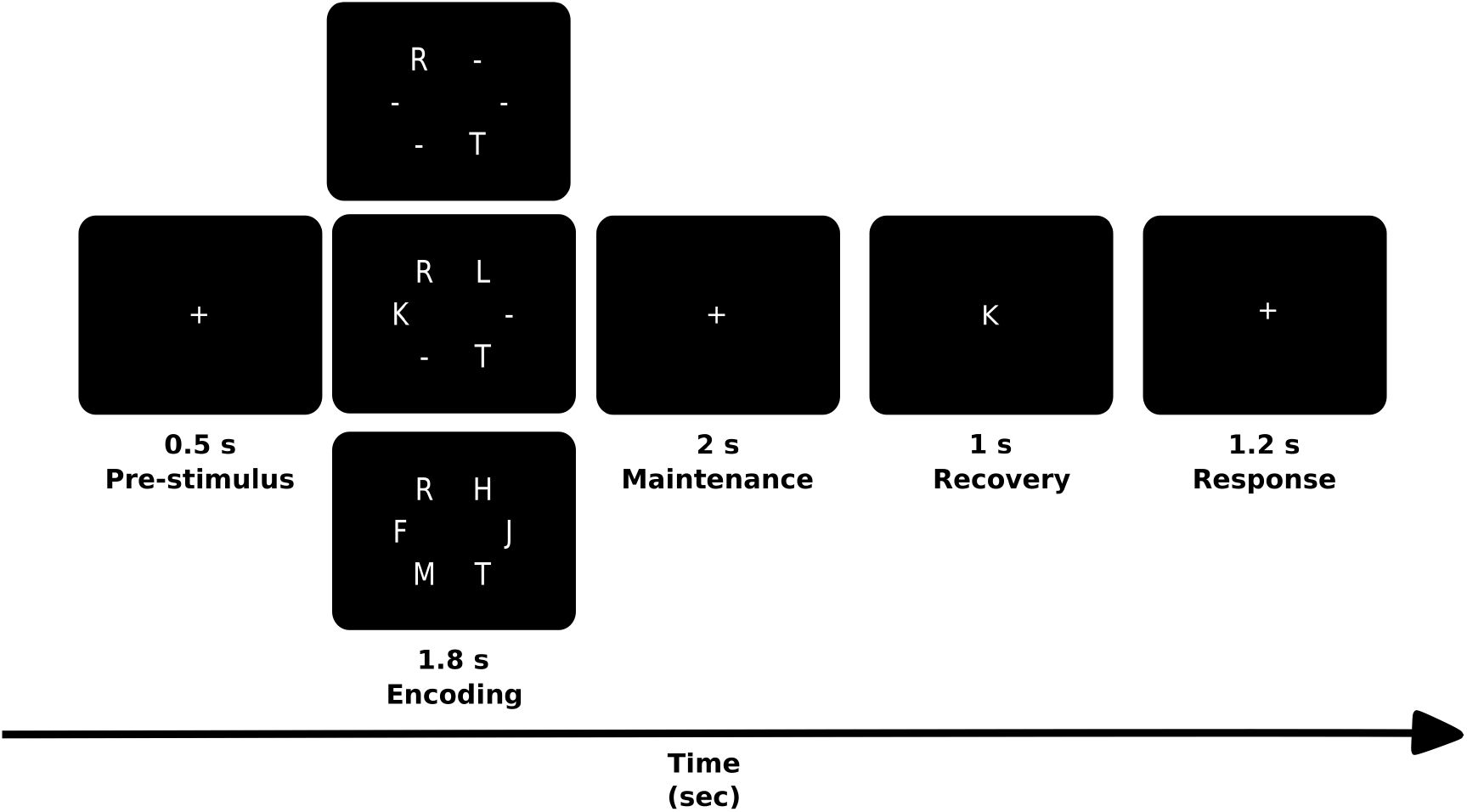
Scheme of the experimental task, adapted from the Sternberg’s Memory Scanning Task described by Jensen et al (2002).

**Table 1.** Demographic and clinical description of the sample. PASAT (Paced Auditory Serial Addition Task) and SDMT (Symbol Digit Modalities Test) evaluates attention, processing speed, and working memory; EDSS (Expanded Disability Status Scale); Brief Visuospatial Memory Test-Revised (BVMT-R) measures visuospatial memory; and the World Health Organization-University of California-Los Angeles Auditory Verbal Learning Test (WHO UCLA AVLT) measures verbal memory.

|  | Patients with Multiple sclerosis (n=20) | Healthy subjects (n=20) |
| --- | --- | --- |
| Age, years | 31.5 (7.34) | 31.1 (8.3) |
| Gender (F/M) | 13/7 | 12/8 |
| Years of Education | 17 (0.63) | 17.25 (1.01) |
| Duration of disease, months | 45.07 (33.75) |  |
| EDSS (score) | 1.0 (0.95) |  |
| PASAT (z-score) | -0.6 (0.88) |  |
| SDMT (z-score) | 0.2 (0.95) |  |
| BVMT-R (z-score) | -0.9 (0.9) |  |
| WHO UCLA AVL-T (z-score) | -0.4 (1.1) |  |
| Stroop (z-score) | 0.4 (0.8) |  |

## Results

### Behavioral

Both groups had over chance performance in all memory load conditions (Wilcoxon test, ps<0.001, Bonferroni corrected), without differences between groups (p>0.2, uncorrected). Patients with multiple sclerosis presented a tendency to have less decrease in their performance in relation to the load increases (difference between load 2 and 6, control mean 0.14, patient mean 0.09, p=0.06, Figure 2). In the following analysis, we focused on the high load memory condition (six items) because there were more errors and there were no differences between groups. We studied the reaction time (RT) as an index of cognitive effort, especially when an error occurred. Both groups presented longer RT for incorrect responses than for correct responses (Wilcoxon test, ps<0.003, Bonferroni corrected) without difference between groups. Additionally, we found that patients presented longer RT for incorrect responses when the probe was part of the memory set (incorrect match responses, EM, Figure 2), which led to significant differences between groups (Wilcoxon tests, p=0.047). Finally, to rule out difference of RT due to motor impairment, we calculated the difference in RT for load memory 2, and the difference in RT using different hands to answer also during load 2. These two measurements were no significantly different between groups (Wilcoxon test, p=0.5, p=0.9 respectively)

**Figure 2.**
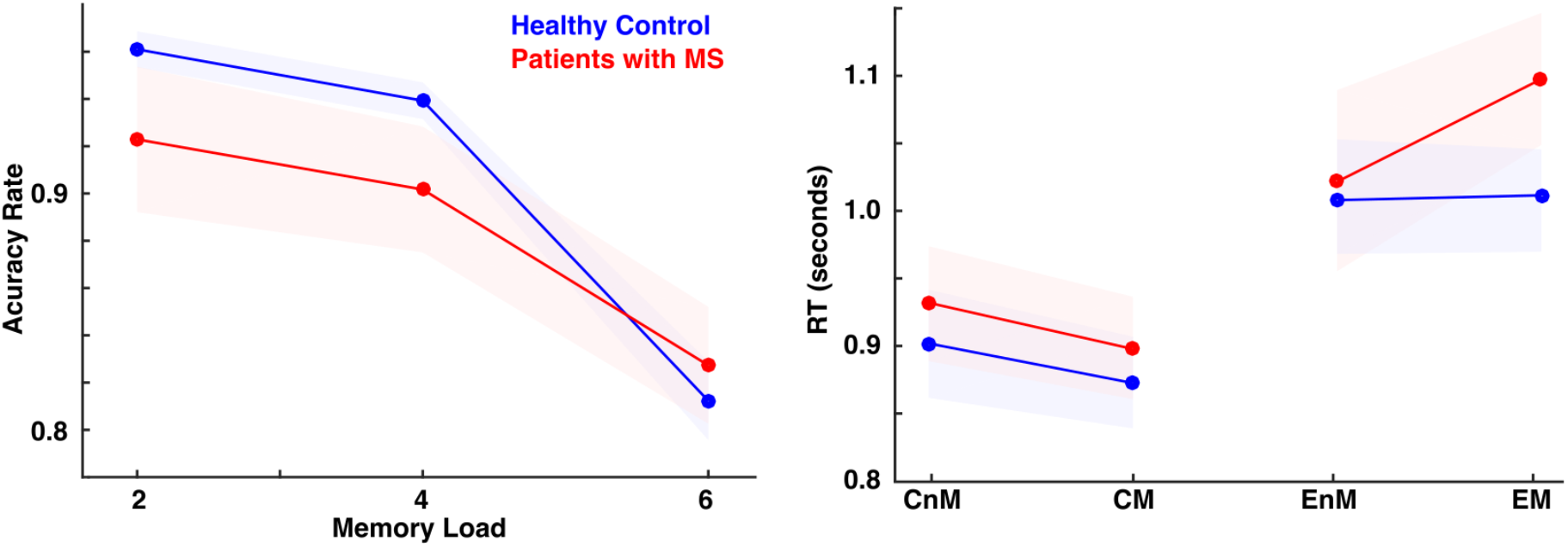
Behavioral Analyses. Left panel shows the Accuracy Rate per memory load. Right panel shows the reaction time (RT) for memory load 6 separated by correct responses for no-matched target (CnM), correct responses for matched target (CM), incorrect responses for no-matched targets (EnM), incorrect responses for matched targets (EM). Red represents patients with multiple sclerosis (MS) and blue healthy control. Colored areas represent standard error of mean.

### Time-Frequency EEG Analysis

We studied the effect of both memory load and successful memory performance on the power of brain oscillatory activity. Regarding the effect of memory load, in the control group we found two effects as expected, a positive modulation in theta activity and a negative modulation in alpha/beta activity ^13,23^. In contrast, we did not find modulations of theta band (6-11 Hz) in patients with multiple sclerosis, leading to a significant difference (p=7 e-7; Cluster Based Permutation (CBP) test) between groups in the initial stage of encoding in this frequency band (between 0.3 and 1 s, Figure 3 D-F). This difference in theta had a topographic distribution located in electrodes of the left hemisphere (Figure 3 I). When analyzing group differences during the maintenance stage, we observed differences in theta activity (5-9 Hz) in the period of 1.3 to 3.3 seconds (p=0.008), which corresponded to the final part of the encoding stage and the entire maintenance phase. The topographic distribution of this modulation was placed over frontoparietal regions with left predominance (Figure 3 M).

**Figure 3.**
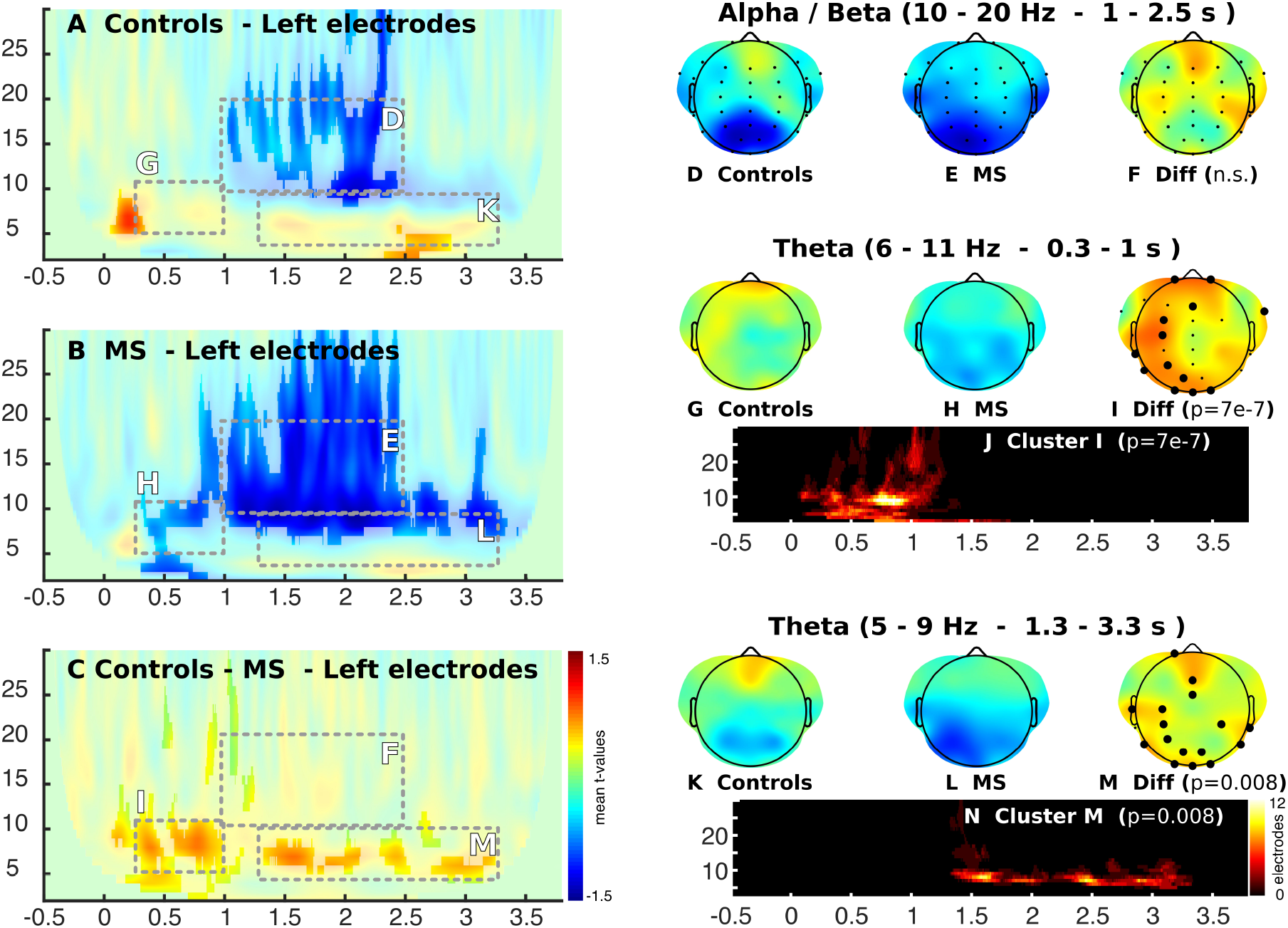
Time-frequency analysis of the effect of memory load (ML) on the difference between multiple sclerosis patients (MS) and healthy control (HC), for the different stages of the adapted Sternberg’s Memory Scanning Task described by Jensen et al (2002). Color represents the mean T-value of the single trial regressions per subjects given by the following equation (Power (f,t) = b1 + b2*ML + b3*SMP). Significant regions are highlighted (CBP test p<0.05).

Next, we analyzed the oscillatory activity related to successful memory performance, that is, the specific activity in the trials in which subjects correctly respond to the target stimuli in relation to those trials where subjects make mistakes (Figure 4). We found that during the maintenance period (2.3 to 2.7 s), patients with multiple sclerosis showed a negative modulation in theta activity (5-11 Hz), while healthy subjects presented a positive modulation (Figure 4A). This led to a significant difference between groups (p=0.01) showing a medial frontal topographic distribution (Fig. 4C). Regarding the analysis of alpha activity (10–15 Hz), we observed that healthy subjects presented a decrease in oscillatory activity in this band in the final period of encoding and beginning of maintenance with an occipital distribution, which was not observed in patients with multiple sclerosis (Figure 4 right). This led to differences between both groups (p=0.008) in parieto-occipital electrodes (Figure 4C right).

**Figure 4.**
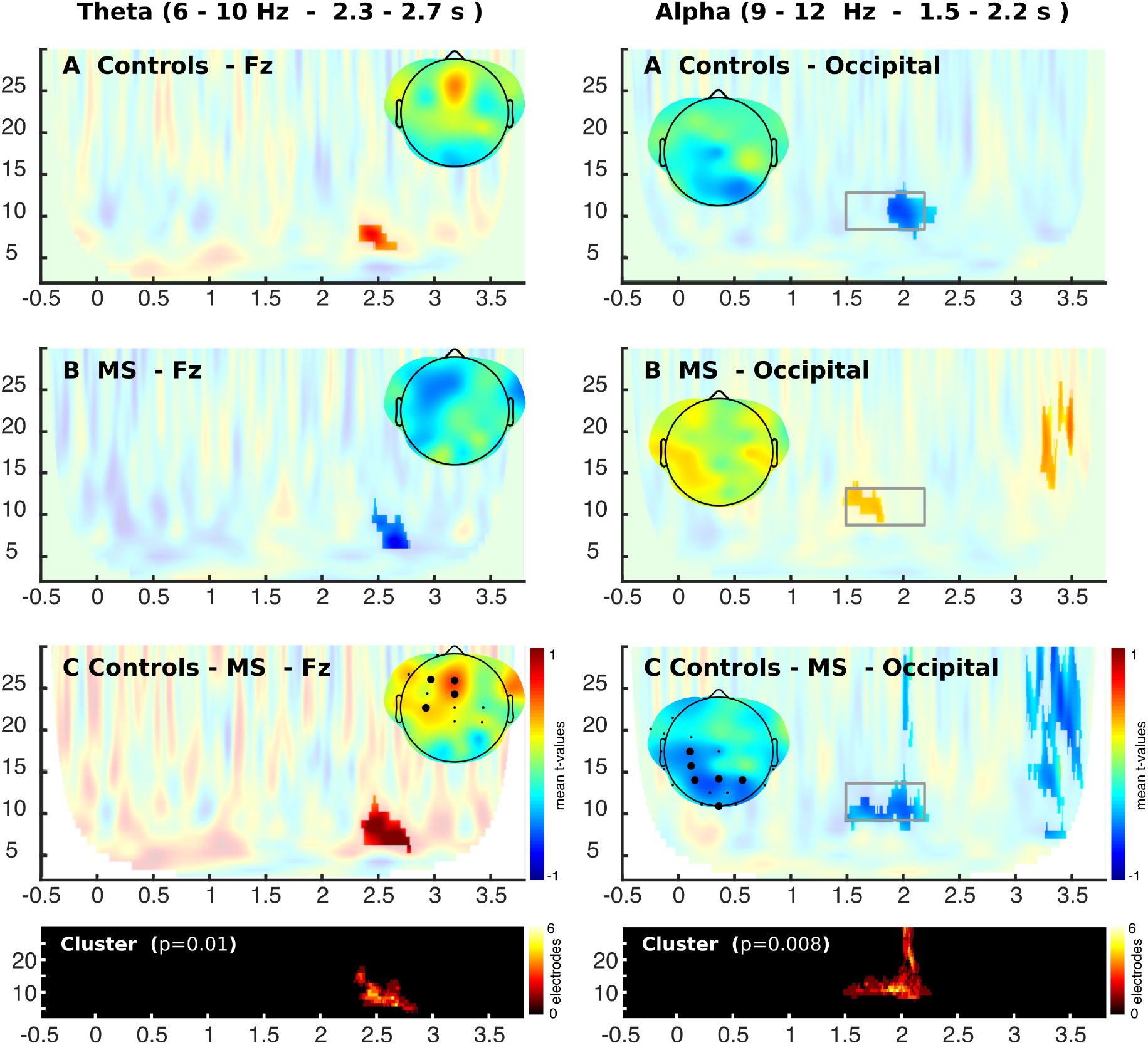
Time-frequency analysis of the effect of successful memory performance (SMP). Colors represent the mean T-value of the single trial regressions per subjects given by the following equation (Power (f,t) = b1 + b2*ML + b3*SMP). Significant regions are highlighted (CBP test p<0.05).

### Source reconstruction

We carried out source reconstructions of the significant differences between groups in theta activity. For encoding we found that theta modulation was placed in left parietal and temporal cortex (Figure 5, False Discovery Rate, FDR, q<0.05 vertex corrected, and p<0.05 cluster-corrected). Two additional clusters were found in right temporal and medial frontal cortex (p<0.05 cluster-corrected). For maintenance, we found that the main modulation was placed again in left temporal and parietal lobes (Figure 5, FDR q<0.05 vertex corrected, and p<0.05 cluster-corrected). Additionally, we found a cluster in right orbitofrontal cortex and right parietal cortex (Figure 5, FDR q<0.05 vertex corrected, and p<0.05 cluster-corrected). For the modulation related to successful memory performance, the source of theta was placed over the medial prefrontal cortex (Figure 5, FDR q<0.05 vertex corrected, and p<0.05 cluster corrected).

**Figure 5.**
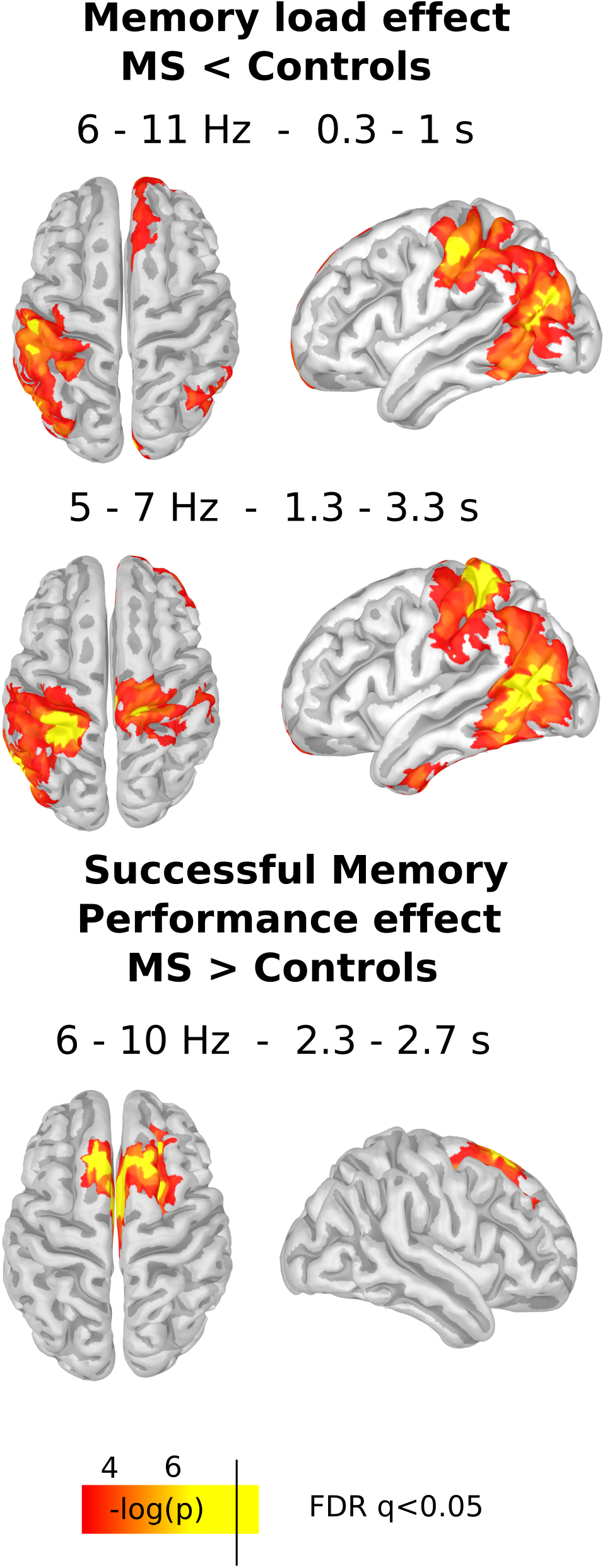
Source reconstruction of the differences between groups. The upper panel shows the differences in the memory load modulation in theta activity during encoding as is highlighted in I. The middle panel shows the differences in the memory load modulation in theta activity during maintenance as is highlighted in Figure 3 M. The top panel shows the differences in the modulation in theta activity related to unsuccessful memory performance during maintenance as is highlighted in Figure 4 C. Only significant clusters (p <0.05 cluster corrected), and vertexes that survive vertex-based correction are shown in yellow (FDR<0.05).

### Connectivity

Considering the results of the oscillatory activity, we carried out a connectivity analysis. We selected a frontal electrode (Fz) and a left parietal electrode (Cp3), since modulation in theta for both memory load and successful memory maintenance were found in these electrodes. The following source reconstruction enabled us to infer that they represent frontal and temporo-parietal activity respectively. We first used GC that measures statistic dependency between signals (e.i., if one signal is useful in forescasting another signal). During the maintenance stage, we found that healthy subjects presented an increase in the parietal-to-frontal connectivity in the time domain, as an indicator of successful memory performance (−0.72, p=0.01). This led to a significant difference in patients with multiple sclerosis, who did not present this modulation (diff, p=0.009, Figure 6 and Table 2). Additionally, this modulation changed in the interaction between memory load and successful memory performance, reversing the direction on high memory loads from frontal-to-parietal (p=0.0089, see Table 2 and Figure 6). Patients with multiple sclerosis did not show this pattern, demonstrating a significant difference when compared to healthy subjects (p=0.04, see Table 2). No significant modulations were found for the encoding period in both groups (Table 2).

**Figure 6.**
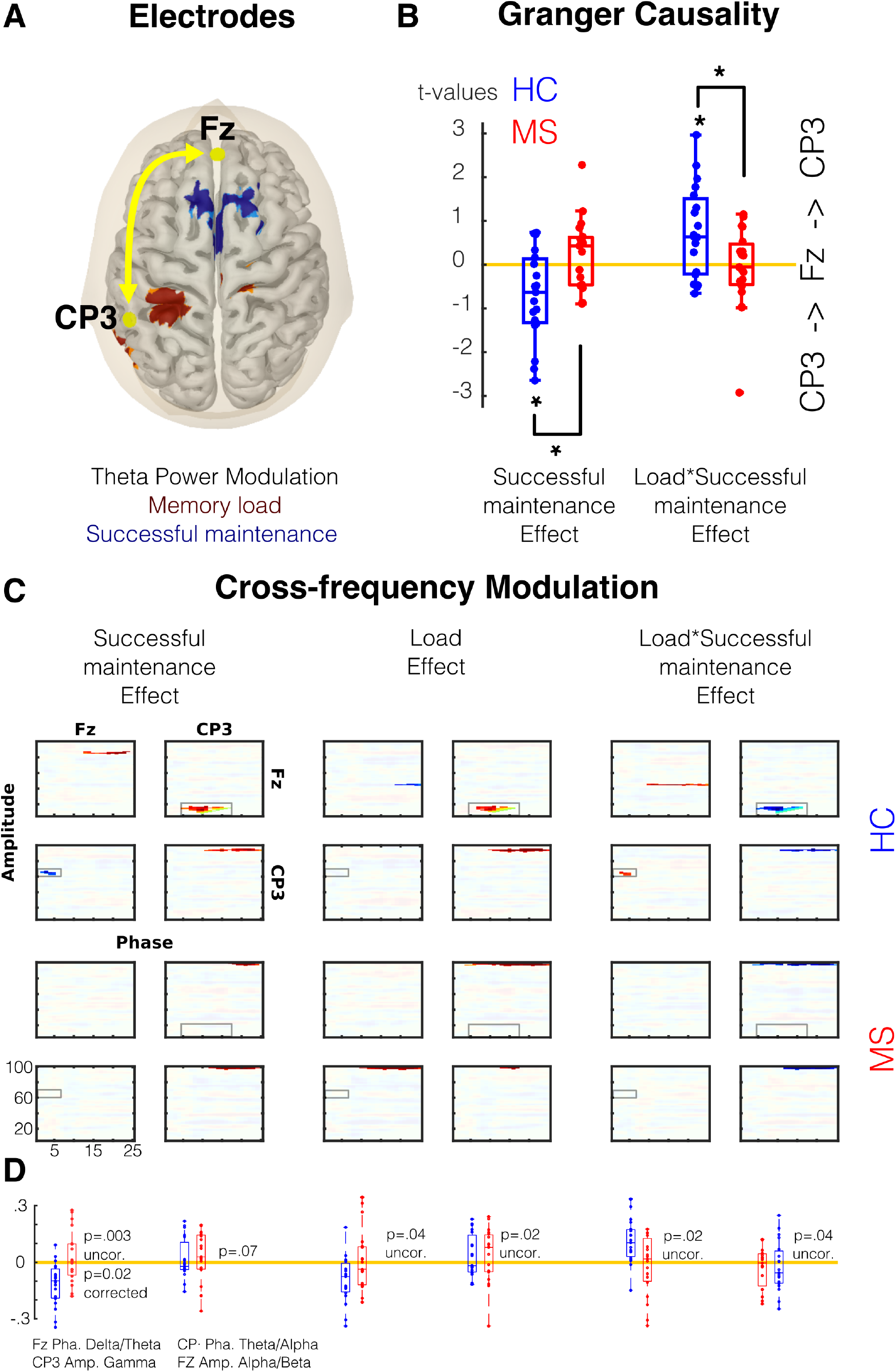
Connectivity Analysis. A. Selected electrode and the source of the theta modulation as showed in Figure 5. B. the t values resulting of single trial models of the Granger Causality between Fz and CP3 electrodes during maintenance (See also table 2.). C Cross-frequency modulation using PAC analysis. Significant areas are highlighted (CBP test p<0.05). D. Comparison between groups in areas where healthy control showed significant modulation. Red depicts Multiple Sclerosis (MS) group and blue Healthy Control (HC) group.

**Table 2.** Single trial regression with both groups Granger Causality between frontal (FZ) and parietal (CP3) electrodes. Positive values indicate frontal to parietal connectivity.

| Frontal to Parietal Connectivity |  |  |  |  |  |  |  |  |  |  |  |
| --- | --- | --- | --- | --- | --- | --- | --- | --- | --- | --- | --- |
| Encoding |  |  |  |  |  | Maintenance |  |  |  |  |  |
|  |  | int | SMP | ML | ML * SMP |  |  | int | SMP | ML | ML * SMP |
| HC |  | -0.54 | -0.31 | 0.23 | 0.24 |  |  | -0.39 | -0.73 | 0.25 | 0.73 |
| p |  | ns | ns | ns | ns |  |  | ns | 0.01 | ns | 0.01 |
| MS |  | 0.3 | 0.36 | -0.2 | -0.34 |  |  | 0.07 | 0.2 | 0.04 | -0.09 |
| p |  | ns | ns | ns | ns |  |  | ns | ns | ns | ns |
| diff |  |  |  |  |  |  |  |  |  |  |  |
| p |  | ns | ns | ns | ns |  |  | ns | 0.01 | ns | 0.04 |
| SMP: Successful memory performance, ML: Memory load, HC: healthy controls, MS: patients with Multiple Sclerosis, ns: non-significant. |  |  |  |  |  |  |  |  |  |  |  |

We then used a complementary approach to infer the flow f the information. Brain connectivity between cortical regions is regulated principally by the projections between supragranular and infragranular layers ^24,25^. Connections originating in superficial and deep layers have distinct spectral fingerprints ^26,27^. Thus, the interactions between frequency bands (analyzed by mean of cross-frequency coupling or PAC) can reveal the information flow ^28,29^. Thus, it is possible to infer that if the phase of a low frequency of one cortical area correlates with the amplitude of a higher frequency of another cortical area, the information flow could have this direction. Using a similar single trial modeling as in GC analyses, we found similar results. In healthy subjects, we found a modulation between the delta/theta phase of frontal area (Fz) and the gamma amplitude of parietal areas (CP3). This modulation correlated negatively with successful memory performance but positively with the interaction between successful memory performance and memory load (Figure 6C). In contrast, we found a second modulation between the theta/alpha phase of parietal areas (CP3) and the beta amplitude of frontal areas (Fz), with the opposite patterns. This second modulation correlated positively with successful memory performance and memory load, but negatively with the interaction between successful memory performance and memory load (Figure 6C). None of these modulations were observed in patients with Multiple Sclerosis, leading to significant differences between groups.

### Clinical Correlations

Finally, we explore whether the frontal to parietal connectivity correlated with clinical assessment. We used the regressor of the interaction between memory load and successful memory performance in Granger Causality analysis and the Neuropsychological test. Interestingly, in the patient group, the frontal-to-parietal connectivity was correlated with PASAT that measure WM capacity (Spearman correlation, rho=0.58, p=0.04 Bonferroni corrected). None of the other tests showed significant correlation (SDMT, rho=0.03, p=0.8; BVMTc, rho=-0.3 p=0.1; CVLT-VII, rho=0.13, p=0.58; Stroop, rho=-0.25, p=0.3; all ps uncorrected).

## Discussion

In this study we assessed the cortical circuits that support verbal working memory in patients with relapsing-remitting multiple sclerosis with minimal or no burden of neurological disability (EDSS <3 and PASAT >-1.5 SD). In the early stage of the disease, patients commonly do not present objective cognitive alterations in the neuropsychological evaluations, however, they manifest a subjective sensation of difficulties in their cognitive performance in daily activities (e.g., occupational or academic tasks) ^30^. These patients report a common clinical pattern of difficulties in the performance of daily cognitive tasks involving working memory, but without clear evidence in the clinical tests applied in their evaluation routines^30–32^. Accordingly, we observed no significant differences in WM performance, but clear neurophysiological differences between patient and control groups. In the early stage of the disease, the clinical evaluations applied to multiple sclerosis patients without impairment often do not detect a deterioration of this cognitive function ^33–35^. The RT difference that we found could reflect an increase in the cognitive effort necessary to correctly solve the task and could be an early behavioral marker of WM impairment. In fact, reaction time analyses are a good marker of attention and cognitive control dynamics ^36–40^.

In spite of no clear evidence for behavioral impairments in the accuracy in the WM task, patients with multiple sclerosis demonstrated a distributed oscillatory activity reorganization. The ability to maintain a sustained activity in the fronto-parietal network in the absence of sensory stimuli depends on the synchronous structured activity of different frequency ranges ^41^. Our working memory task elicited a recognized oscillatory activity in a frontal and parietal network ^11,42–44^. The left lateralization of the power of the theta oscillatory activity could be related to the areas involved in language, i.e., the phonological loop, required for the specific information of our stimuli (consonants) ^13,45^. Low frequency synchronization between the medial frontal region and temporal cortex has been demonstrated in non-verbal WM tasks, probably reflecting cognitive control ^15,23^. Indeed, our connectivity analyses showed specific dynamics, in which frontal to parietal/temporal influences increased in function of memory load when a subsequent successful memory performance occurs. These findings are in line with several recent reports indicating that frontal interactions with other cortical areas are key aspects of successful memory performance ^14,46,47^.

In contrast, patients with multiple sclerosis demonstrated a loss of the WM oscillatory dynamics. These patterns could represent an inefficient cognitive effort to keep the stimuli information active in WM, which would be in accordance with the longer reaction times for erroneous responses in high memory load. In addition to this, the patient group did not present an increase in frontal-to-parietal connectivity ^48,49^. WM requires the synchronization of neural network connections distributed in the prefrontal and parietal regions in order to integrate complex information for generating appropriate responses. These distributed brain networks are especially sensitive to the diffuse damage of white and gray substance found in multiple sclerosis ^17,18,50^.

Thus, our results are in accordance with recent findings in patients with mild or minimal cognitive deficit that show reorganization in electrophysiological activity ^49,51^. In spite of the fact that patients could have fewer resources to maintain the stimuli in WM they can achieve a similar performance to healthy subjects, as long as the tasks are not too, by means of redistribution of the remaining and available resources. It has been proposed that the central executive may be the main component of WM that is disrupted in cognitively impaired multiple sclerosis patients^52^. This proposition is supported indirectly by evidence that suggests that the dorsolateral prefrontal cortex, which is thought to underlie executive control, is commonly recruited when there is heavy demand placed on WM in individuals with brain injury ^53^. Our findings give further evidence to support this interpretation. Indeed, the frontal low frequency influences the gamma power in parietal and temporal areas, and this influence increases in relation to memory load increase. Patients fail to display this influence, and this failure could reflect the loss of the mechanisms by which the control process carried out by frontal areas produces a successful memory performance.

Studies using fMRI have found several changes in brain activity during WM tasks including decrease and increase in both connectivity and activity ^54,55^. Indeed, follow-up studies show an inverted-U form in the evolution of the disease in resting-state functional connectivity^56^. Early changes in the connectivity/functional patterns have been interpreted as compensatory changes in prefrontal cortical regions underlying modulations of executive aspects of WM ^57–59^. For instance, studies in very early states of multiple sclerosis have found medial PFC activity increases, although later meta-analyses studies have revealed a decrease in medial prefrontal activity. In this context, our study gives several insights into the participation of medial prefrontal activity in working memory and its relationship with the cognitive dysfunction in patients with multiple sclerosis. Medial prefrontal theta activity is related specifically to subsequent successful memory performances. As this oscillatory activity has been generally related to cognitive control in several tasks ^60^, it is possible to interpret this activity index as the cognitive effort related to successful performance. Interestingly, patients present more activity for errors and also present an increase of reaction time for incorrect responses. Thus, the oscillatory pattern demonstrated by multiple sclerosis patients may indicate a decrease in the detection of the need for additional cognitive control ^61,62^. In line with this, patients do not present an increase of frontal to parietal influences in relation to successful memory. Indeed, patients with Multiple Sclerosis and cognitive impairments demonstrate low frequency amplitude decreases during cognitive tasks ^63^. Along with this, it has been observed that the nested frequencies between theta and gamma can represent the way in which multiple maintenance items are organized into WM ^41^. The nested frequencies or PAC refers to a specific type of oscillatory activity, which would reflect a general mechanism by which the cortical areas organize and structure information ^64–67^. Accordingly, we observed a modulation between delta/theta and gamma frequencies in healthy subjects. This communication channel seems specific to the frontal to parietal communication and shows an increase in relation to memory load. While, for low memory load, in information flow seem to be predominantly from parietal to frontal, using a different nested frequency. Patients did not demonstrate these dynamics, reflecting the poor or inefficient frontal parietal communication. Interestingly, this loss of interplay between the frontal and parietal oscillatory dynamics could be an early marker of working memory deficits in multiple sclerosis patients. Thus, these patients’ performance suggests an impairment in the establishment and maintenance of a fluid oscillatory dialogue between the areas involved in the successful development of a working memory task. Indeed, connectivity pattern correlated with clinical test of WM.

In summary, in spite of a high incidence of cognitive deficits multiple sclerosis and a large body of literature investigating cognitive dysfunction, the specific oscillatory features that drive cognitive deterioration in multiple sclerosis remain unclear. Hence, our results demonstrate a specific electrophysiological mechanism underlying the WM deficit in Patients with Multiple Sclerosis. Several investigations have revealed that cerebral oscillatory activity supports different cognitive processes and have indexed their alteration in clinical populations ^68–70^. Indeed, recent evidence indicates that increasing theta activity by means of transcranial Alternating Current Stimulation can improve WM performance in healthy and aging subjects ^14,47^. It has been seen that cognitive training can increase memory capacity and that this increase correlates with changes in connectivity of distant brain areas, specifically between frontal and parietal regions. Therefore, the specific oscillatory features related to WM deficits identified in our study could serve to implement non-pharmacological treatments using non-invasive brain stimulation and cognitive training, in order to contribute to the improvement of the quality of life of these patients.

## Methods

### Design and Participants

Our study is a case-control design that include 40 participants. A sample consisted of 20 patients with relapsing-remitting multiple sclerosis in an early stage, with minimal to no clinical evidence of cognitive alterations (Table 1). According to the 2010 McDonald Criteria, the medical diagnosis was made by a Neurologist ^71^. Stable patients without episodes of relapses in the last month, with scores of three or more on the Expanded Disability Status Scale (EDSS), with less than −1.5 z-score of Paced Auditory Serial Addition Test (PASAT), with non-correctable visual alterations, with a history of traumatic brain injury, neurological and/or psychiatric pathologies, and abuse or regular consumption of drugs or alcohol were excluded. The patients participated in the study while on their usual disease modifying therapies (i.e., immunomodulation therapy only, without other treatment, such as antidepressants or Benzodiazepines). The control group was composed of 20 healthy volunteers, comparable in age, sex, manual preference, and educational level (Table 1). As well as in the patient group, healthy subjects with non-correctable visual alterations, with a history of neurological and/or psychiatric pathologies, traumatic brain injury, and abuse or regular consumption of drugs or alcohol were excluded. All participants were Spanish native speakers and provided signed informed consent prior to participation in the study. Patients underwent neuropsychological assessment during the month previous to the EEG session. This was their first neuropsychological evaluation. These assessments included PASAT that measures cognitive processing speed and working memory, Symbol Digit Modalities Test (SDMT) that measures cognitive processing speed, the Brief Visuospatial Memory Test-Revised (BVMT-R) that measures visuospatial memory, and the World Health Organization-University of California-Los Angeles Auditory Verbal Learning Test (WHO UCLA AVLT) that measures verbal memory. Details are summarized in Table 1.

The experimental protocol and all methods were performance in accordance to institutional guidelines and were approved by the Ethical Committee of the Pontificia Universidad Católica de Chile.

### Sample size

For the estimation of the minimum required sample size the following parameters were considered: a) Effect size for the mixed ANOVA statistical test (2 x 2, with interaction effects), b) Statistical power (1-β)=.95 and c) Significance level **α**=.05. Considering an effect size *η*2 =0.09 (effect size F = 0.3 ^13,72^), the sample size amounts to a total of 40 participants (n1=20; n2=20).

### Experimental Task

In this study we implement a modified version of Sternberg’s Memory Scanning task [Jensen, 2002]. This task consisted of a list of consonants simultaneously presented and displayed in a circular arrangement with a fixation cross in the center of a computer monitor located 57 cm from the subject. The letter “Y” was not included in the memory set to avoid the generation of words that could be used as clues by the subjects. Each memory set arrangement consisted of groups of two, four, or six consonants generating three levels of WM load. The latter refers to the progressive number of stimuli to be stored and manipulated in WM. All the stimuli were placed foveally, minimizing the effect generated by saccadic movements (see the experimental task outlined in Figure 1). Each memory set was presented for 1800 ms (encoding period), followed by a black screen with the fixation cross (maintenance period) of 2000 ms and then a recovery period in which the fixation point was replaced by a target stimulus for 1000 ms. The subjects were instructed to memorize the memory set and then report whether the target stimulus was present or absent in the memory set, using the right or left hand alternately. Subjects had 2200 ms to answer. Each subject had to respond 270 trials (90 trials for each memory load set). The trials were presented in two main blocks divided by a pause regulated by the subject. In addition, each main block was constituted by 15 sub-blocks, formed by 9 trials of the same memory load each. The order of the memory load was randomized. Both the presentation of the stimuli and the recording of the test responses were done with the Software Presentation^®^ (Version 13.0, www.neurobs.com).

### Electrophysiological Recordings

Continuous EEG recordings were obtained with a 40-electrode EEG System (NuAmps, Neuroscan). All impedances were kept under 5kΩ. Electrode impedance was retested during pauses to ensure stable values throughout the experiment. All electrodes were referenced to averaged mastoids during acquisition and the signal was digitized at 1 kHz. Electro-oculogram was obtained with four electrodes. All recordings were acquired using Scan 4.3 and stored for off-line analysis. At the end of each session, electrode position and head points were digitalized using a 3D tracking system (Polhemus Isotrak).

### Electrophysiological data analysis

EEG signals were preprocessed using a 0.1-100 Hz band-pass filter. Eye blinks were identified by a threshold criterion of ±100 μV, and their contribution was removed from each dataset using Independent Component Analysis (ICA). Other remaining artifacts (e.g., muscular artifacts) were detected by visual inspection of both the raw signal and the spectrogram. We thus obtained 243 ± 28 artifact-free trials per subject. All artifact-free trials were transformed into current source density (CSD) that was estimated using the spherical spline surface Laplacian algorithm suggested by Perrin et al.,^73^ and implemented by Kayser and Tenke ^74,75^. Induced power distribution was computed using Wavelet transform, with a 5-cycle Morlet wavelet, in a −0.5 to 3.8 s window around the onset of the memory set stimuli. This timewindow includes 0.5 seconds of inter stimulus interval, 1.8 seconds of the stimulus of the memory set and 2 seconds of maintenance period. For all analyses, we used the dB of power related to the baseline (15 seconds acquired in the beginning of each block).

### Source Reconstruction

The neural current density time series at source levels were calculated by applying a weighted minimum norm to estimate inverse solution ^76^ with unconstrained dipole orientations in single trials as in prior work^68,77,78^. We used a default anatomy of the Montreal Neurological Institute (MNI/Colin27) wrapped to the individual head shape (using ~300 head points per subject). We defined 3 x 4000 sources constrained to the segmented gray cortical volume (3 orthogonal sources at each spatial location) in order to compute a three-layer (scalp, inner skull, outer skull) boundary element conductivity model and the physical forward model ^79^. Since the inverse solution is a linear transformation, it does not modify the spectral content of the underlying sources. Therefore, it is possible to undertake time-frequency analyses directly in the source space. Finally, we reduced the number of sources by keeping a single source at each spatial location that pointed into the direction of maximal variance. For this, we applied a principal component analysis to the covariance matrix obtained from the 3 orthogonal time series estimated at each source location. Since we used a small number of electrodes (40) and no individual anatomy for head model calculation, the spatial precision of the source estimations is limited. In order to minimize the possibility of erroneous results, we only present source estimations if there are both statistically significant differences at the electrode level and the differences at the source levels survive a multiple comparison correction (cluster-based permutation test and vertex correction using false discovery rate, q=0.05).

### Statistical analysis

We used the Kolmogorov-Smirnoff to test for normality. When the data did not meet the normal assumption, we used non-parametric tests. We evaluated pair comparisons using Wilcoxon test and Bonferroni correction. For the EEG statistical analysis, we first fitted a General Linear Model (GLM) of the power of the oscillatory activity per trial in each subject (first level analysis, see ^13,70,80^),

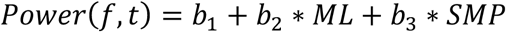

where *b*_1_ is the intercept, and *b*_2_ is the slope or coefficient for the variable Memory Load (ML, ordinal variable that takes the value 2, 4, or 6 depending on the memory load condition) and *b*_3_ is the slope for the variable Successful Memory Performance (SMP, dummy variable that takes the value 0 if the subject makes a mistake in this trail or 1 if the target stimulus is correctly identified). We thus obtained a 3D matrix of t-values (sensor, time, frequency) for each regressor and subject. We then explored for differences between groups and conditions using the Wilcoxon test (second level analysis). To correct for multiple comparisons in timefrequency charts, we used the Cluster-based Permutation (CBP) test ^81^ using 1000 permutations. For more detail see prior work ^82,83^.

### Causal interactions

To evaluate the influence of frontal regions over temporo-parietal regions, we estimated Granger Causality (GC) ^84^ between selected electrodes (Fz and CP3). The causality was calculated over time series per trials. See detail in^83,84^. We obtained a GC term per trial that was then used in the modeling analyses (see below).

### Phase-amplitude coupling (PAC)

To evaluate cross-frequency modulation, we carried out a phase-amplitude coupling analysis (PAC) as described in ^64^. Briefly, for a given frequency pair, the raw signal was filtered separately in both frequencies (zero phase shift non-causal finite impulse filter with 0.5 Hz roll-off). The lower frequency ranged from 1 to 35 Hz (0.4 Hz increments, 0.8 Hz filtered bandwidth) and the higher ranged from 5 to 120 Hz (1 Hz increments, 5 Hz filtered bandwidth). The phase of the lower frequency range and the amplitude of the higher frequency range were computed using Hilbert transformation. For each epoch of maintenance per trial (1.8 to 3.8 s), we computed the circular-linear correlation between the phase of lower frequencies and the amplitude of higher frequencies. Thus, we obtained a circular correlation coefficient (CCC) per trial and per combination of pairs between the selected electrodes (Fz-Fz, Fz-Cp3, Cp3-Fz, Cp3-Cp3). Thus, we obtained a CCC per trial and pair of electrodes that were used in the modeling analyses (see below).

### GC and PAC modeling

For both GC and PAC analyses, we used the following modeling. At subject level analysis (first level analysis), we compared if the circular correlation coefficient (CCC) or granger coefficient (GC) variation correlated with the task parameters using multiple linear regression as the following equations depict:

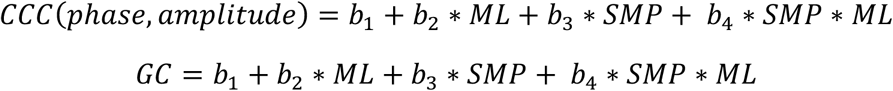

At group level analysis per frequency pair, we compared whether t values of each regressor were statistically different from zero using the Wilcoxon signed-rank test. For PAC analyses, we corrected for multiple comparisons using a CBP test. The initial threshold for cluster detection was p<0.05, and the final threshold for significant cluster was p<0.01. For between-group comparison we used Wilcoxon rank-sum test and Bonferroni correction.

### Software

All behavioral statistical analyses were performed in R. The EEG signal processing was implemented in MATLAB using CSD toolbox, in-house scripts (available online as LANtoolboxhttp://neuroCICS.udd.cl/LANtoolbox.html), BrainStorm ^85^ and open MEEG toolboxes ^86^.

## Data availability

All Data and script used for data analyses are freely available on http://neuroCICS.udd.cl

## Author contributions

A.F-V., C.C., F.Z., F.A., P.B. designed the experiment; P.B., F.Z. programmed the experiment; A.F-V., F.Z., P.B. conducted the experiments; C.C., E.C., R.U., M.V. evaluated the participants; A.F-V., R.H-C., P.B. analyzed the data; A.F-V., R.H-C., P.B., F.Z., F.A. interpreted and discussed the data; A.F-V., R.H-C., F.A., P.B. wrote the manuscript.

